# Consistent individual positions within roosts in Spix’s disc-winged bats

**DOI:** 10.1101/2022.11.04.515223

**Authors:** Giada Giacomini, Silvia Chaves-Ramírez, H. Andrés Hernández-Pinsón, José Pablo Barrantes, Gloriana Chaverri

**Affiliations:** Research Centre in Evolutionary Anthropology and Palaeoecology, School of Biological and Environmental Sciences, Liverpool John Moores University, UK; Programa de Posgrado en Biología, Universidad de Costa Rica, San José, Costa Rica; Programa de Posgrado en Computación e Informática, Universidad de Costa Rica, San José, Costa Rica; Sede del Sur, Universidad de Costa Rica, Golfito, Costa Rica; Smithsonian Tropical Research Institute, Balboa, Ancón, República de Panamá

**Keywords:** Bats, dominance, linearity, Resource Holding Potential, roosts

## Abstract

Individuals within both moving and stationary groups arrange themselves in a predictable manner; for example, some individuals are consistently found at the front of the group or in the periphery and others in the center. Each position may be associated with various costs, such as greater exposure to predators, and benefits, such as preferential access to food. In social bats, we would expect a similar consistent arrangement for groups at roost-sites, which is where these mammals spend the largest portion of their lives. Here we study the relative position of individuals within a roost-site and establish if sex, age, and vocal behavior are associated with a given position. We focus on the highly cohesive and mobile social groups found in Spix’s disc-winged bats (*Thyroptera tricolor*) given this species’ use of a tubular roosting structure that forces individuals to be arranged linearly within its internal space. We obtained high scores for linearity measures, particularly for the top and bottom positions, indicating that bats position themselves in a predictable way despite constant roost-switching. We also found that sex and age were associated with the use of certain positions within the roost; for example, males and subadults tend to occupy the top part (near the roost’s entrance) more often than expected by chance. Our results demonstrate, for the first time, that bats are capable of maintaining a consistent and predictable position within their roosts despite having to relocate daily, and that there is a link between individual traits and position preferences.

## INTRODUCTION

The relative spatial position of individuals within groups is relevant for understanding the consequences of social living, as spatial positions may determine vulnerability to predation and access to resources (Bumann et al., 1997; Morrell & Romey, 2008). For example, animals positioned at the front of a moving group need to invest more time in vigilance and are also subject to significant predation risks (van Schaik & van Noordwijk, 1989; Dirk et al., 1997; Di Blanco & Hirsch, 2006), but they may also have greater feeding success (Hirsch, 2011). For stationary groups, studies have shown that individuals are often positioned predictably either on the periphery or the center of the group, and that the former are more vulnerable to predation (Rayor & Uetz, 1990; McGowan et al., 2006). What is common to both moving and stationary groups is that these risks and costs are highly influenced by site-specific conditions; for example, the risks faced or perceived by individuals in the periphery of groups may vary depending on a predator’s attack strategy and distance, in addition to habitat structure (Hirsch & Morrell, 2011; Thaker et al., 2011; Minias, 2014; Jolles et al., 2022).

Many animals spend a large portion of their lives within structures that they use for roosting (Ellerman, 1956; Kunz, 1982; Übernickel et al., 2021). In these sites, individuals can avoid adverse environmental conditions and predators; they also conduct many important fitness-related activities, such as grooming, feeding, nursing, and copulating (Hudson & Distel, 1982; Kunz, 1982; Long, 2009; Suselbeek et al., 2014). Roosts have heterogeneous internal conditions (Burda et al., 2007; Ficetola et al., 2012), and thus protection against predators and/or inclement weather is not equally distributed within them. Thus, individuals sharing the same roost most likely scramble to occupy the best positions, as has been observed for stationary groups in many taxa. In colonial spiders (*Metepeira incrassate*; Rayor & Uetz, 1990), starlings (*Sturnus vulgaris*; Summers et al., 1987), and long-tailed tits (*Aegithalos caudatus*; McGowan et al., 2006), some individuals are typically positioned in the center of the group or colony, and monopolization of these preferred locations is based on individuals’ dominance status. However, despite the importance of roosts for the lives of many organisms, and the accumulated evidence of consistent differences in individual positions within both moving and stationary groups, there are only a few studies that have addressed whether animals consistently select specific positions within their colonies while roosting.

Here we test whether Spix’s disc-winged bats (*Thyroptera tricolor*) consistently select specific positions within their groups while using diurnal roosts. This nocturnal bat species lives in highly cohesive and relatively small groups of approximately 5-6 individuals (Chaverri & Gillam, 2016). We take advantage of two unique aspects of the natural history of *T. tricolor*: its use of a tubular roosting structure that seems to force groups to be arranged linearly within its internal space (Findley & Wilson, 1974), and the need to relocate daily to a new roost site. The latter occurs because the tubular shape of furled leaves of plants in the order Zingiberales (e.g., *Heliconia*, *Calathea*, *Musa*) used as roost-sites unfurl in approximately 24 hours (Vonhof & Fenton, 2004), which presumably forces group members to scramble for the best position within the new tube on a daily basis. While we provide no assessment of costs and benefits of different locations within the leaf’s tubular structure, we explore how certain traits may be associated to specific positions. This will allow us to provide initial clues as to whether some form of dominance relationships mediate the selection of specific positions and develop new hypotheses to test in further studies.

In societies with linear hierarchies, it is often observed that a certain set of individuals dominates over other group members and thus can monopolize resources or ideal positions (Herrera & Macdonald, 1993; Wittig & Boesch, 2003). This effect is referred to as resource holding potential (RHP; (Parker, 1974; Taylor & Elwood, 2003). In many taxa, RHP is associated with body size (Holand et al., 2004; Roden et al., 2005); therefore, if there is sexual dimorphism in that trait, then a possibility is that the larger sex will also have greater RHP. Age is also associated with RHP, since adults typically dominate over juveniles or subadults (Holand et al., 2004; Roden et al., 2005; Lu et al., 2013). Therefore, in our study we predict that females, which are larger than males (Chaverri & Vonhof, 2011), and adults will preferentially occupy certain positions within the roost. Since we have not assessed the costs and benefits of different positions within the roost, we cannot accurately predict which position will be primarily occupied by individuals with greater RHP. However, previous studies on colonially breeding birds show higher consistency of individuals occupying outer positions compared to inner ones, most likely since dominant individuals are constantly fighting for access to central positions while subordinate individuals are consistently relegated to the periphery (McGowan et al., 2006). Thus, we can predict that peripheral positions will be more stable while central ones will be more labile. Also, we predict that the bottom position will be primarily occupied by vocal bats, as previous studies suggest that the most vocal individuals are also the ones that first discover the roost site (Sagot et al., 2018). We assess vocal behavior based on whether individuals produce (vocal) or not produce (nonvocal) response calls, which are emitted by bats that have found a suitable roost site (Chaverri et al., 2010), and those calls promptly recruit other group members which presumably would position themselves above the bat that first located the roost.

## METHODS

### Data acquisition

We collected data on *T. tricolor*’s roosting behavior between November 2017 and mid-January 2018 at Barú Biological Station in southwestern Costa Rica. During this period, most females are in the early stages of gestation; parturition first occurs in February (Chaverri & Vonhof, 2011). We consistently searched the “Pizote” and “Vuelta del Zahíno” trails, an area of approximately 8 ha where we had previously found groups of bats, for a total of 41 days. Each day we searched for furled leaves to identify potential roosts, i.e., leaves which are occupied by a set of individuals (between 1-11, average = 5; Chaverri & Gillam, 2016); bats found in the same roost simultaneously are considered to belong to the same social group. The population of *T. tricolor* in this area has been thoroughly studied (Chaverri et al., 2020) and many bats carry a PIT tag (Biomark Inc., Boise, USA) implanted subcutaneously in the mid-dorsal area. Each tag corresponds to a unique alphanumeric code and allows individual-level identification. We trapped bats by closing the entrance of the tubular leaf used as a roost-site and directing all individuals into a plastic bag and then transferred them to a cloth bag. We installed PIT tags when missing (permit SINAC-ACOPAC-RES-INV-008-2017) and recorded the sex and age of each bat. A few days (range = 1 to 9, mean = 3.8, SD = 2.8) after capturing all individuals in a group, which are highly stable over long periods of time (Chaverri, 2010), we recorded the relative position of each individual within the tubular structure (**Figure 1**) by carefully scanning the leaf from top to bottom with an HPR Plus reader and custom-made antenna (Biomark Inc., Boise, USA). We collected data from all bats found within our sampling area, which comprise a total of 18 different social groups. For analyses we only considered groups that were observed on 5 or more days.

**Figure 1.**
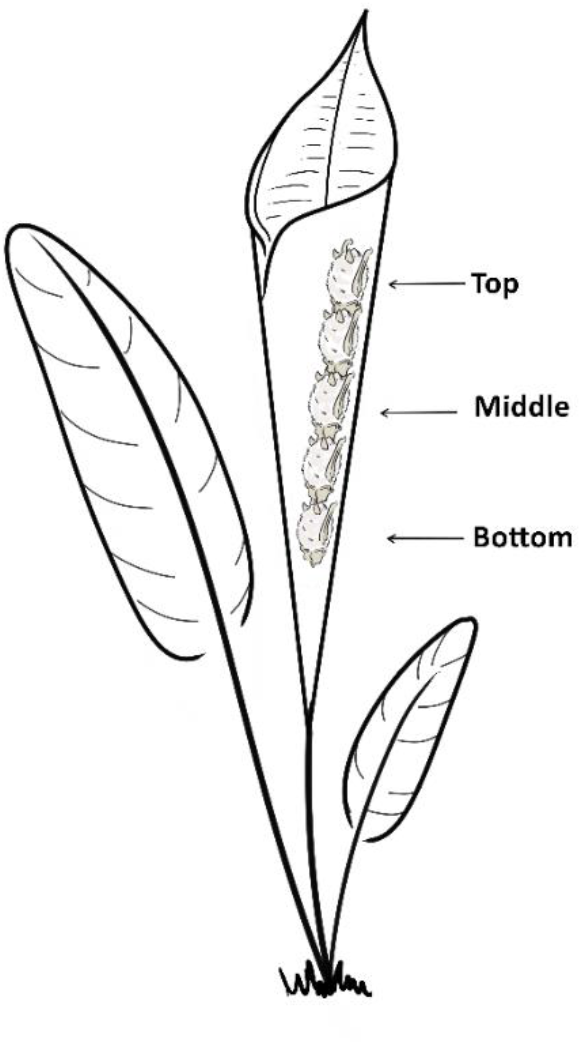
Drawing that shows how bats position themselves within the tubular leaf while roosting, and which individual was located at the top, middle, and bottom. Illustrated by Silvia Chaves-Ramírez.

To test if vocal behavior is correlated to roosting position, we also analyzed the maximum number of response calls emitted by the same identified bats following methods used in previous studies (Chaverri et al., 2020). For this, we placed individual bats within a tubular structure while broadcasting inquiry calls, which prompt the emission of response calls (Chaverri et al., 2010), using a broadband speaker (Ultrasonic Omnidirectional Dynamic Speaker Vifa, Avisoft Bioacoustics, Glienicke/Nordbahn, Germany) connected to an Avisoft UltraSoundGate Player. We recorded response calls using an Avisoft condenser microphone (CM16) through Avisoft’s UltraSoundGate 116Hm connected to a laptop computer running Avisoft-Recorder software (sampling rate 500 kHz, 16-bit resolution). Recording sessions lasted for 1, 5 or 10 minutes (Chaverri et al., 2020). We analyzed recordings using SASLab Pro (Avisoft Bioacoustics).

### Linearity and dominance

We were interested in estimating whether bats consistently positioned themselves in a specific order within the tubular leaf and thus obtained two measures: linearity and dominance. Linearity in our study would mean that, for example, bat *a* is always in the bottom, bat *b* always goes second, bat *c* always goes third, and so on. Linearity was estimated using de Vries’ (de Vries, 1995) *h’*, a modification of Landau’s (1951) h, which measures the level of consistency in linearity and goes from 0 (each individual dominates exactly half the others) to 1 (a perfectly linear hierarchy). As we had no a priori expectation of which position would be dominant, we estimated linearity for all possible arrangements within the tubular structure: top to bottom, middle to bottom/top, and bottom to top. This analysis produces one linearity estimate per group per position.

We verified the consistency in the selection of relative position within a roost by first quantifying the proportion of times each individual was located at each position. Then we randomized the location of individuals, while maintaining group and date consistent, 1000 times and compared the original to the randomized proportions to determine whether individuals use the same location repeatedly or whether roost positions are random. We also calculated which individual(s), per group, were consistently found in a specific position within the tubular structure. For this we estimated David’s (1987) score for each group member. The score produces large positive values for individuals that typically “dominate” (they are consistently found in the top, the middle, or the bottom) and large negative values for individuals that are “dominated”. However, note that we are not measuring dominance per se, that is, the relationship between two individuals which can be explained by one individual (the subordinate) submitting to another (dominant) individual in contest situations (Kaufmann, 1983), but rather the tendency of a given bat to be primarily located in a given position, compared to other group members, within the tubular structure. Again, as we had no a priori expectation of which position should be preferred (or “dominated”), we calculated three David’s scores per individual: one assuming that the individual at the top is dominant, one where the middle is dominant, and one where the bottom is dominant.

To calculate linearity and “dominance” indices, we input the data in SOCPROG (Whitehead, 2009) in group mode allowing for interactions between all group members and for asymmetric interactions within the group. We organized the input data by collection date for each group with the unique bat codes ordered in each entry line. The software assumes that if bat *a* is the first of the sequence, it “dominates” over the other bats. Therefore, we arranged the data in three different formats in order to compute the indices considering 1) the top position as dominant (bats in order from top to bottom: *a, b, c, d, e*); 2) the bottom position as dominant (bats in order from the bottom to the top: *e, d, c, b, a*); 3) and the middle position as dominant (bats in order from the middle to the two extremes: *c, d* & *b* in random order, *e* & *a* in random order). For each group, we analyzed the three datasets separately.

### Statistical analysis

To determine which position, bottom, middle or top, was associated to a more consistent placement of bats, we first compared variation in linearity values among positions using a Levene’s test. The results of the Levene’s test were significant (F_2,27_ = 4.62, P = 0.01), indicating differences in the variation of linearity values among positions. Given the latter, we used a Welch’s ANOVA to test if there were differences in the values of linearity among positions, applying pairwise t-tests, with no assumption of equal variances, to compare linearity estimates among positions.

We determined whether sex, age, and vocal behavior affected “dominance” (David’s score) at particular positions per group within the tubular structure. For this we only considered the individuals with the highest score within each group per position. This typically resulted in a data set comprised of 10 values, 1 per group (see exceptions below). To these 10 values, and for each explanatory variable separately (i.e., sex, age, and vocal behavior), we applied Bayesian binomial tests in the BayesianFirstAid R package (Bååth, 2014) to determine the probability of successes, i.e., instances in which females, adults or vocal bats “dominated” a given position (see predictions). Binomial tests are used to determine whether a proportion of observed values in a binary variable is equal to a hypothesized proportion. The Bayesian binomial test further incorporates subjective prior beliefs about the parameter of interest and updates them with observed data to obtain a posterior probability distribution (Bååth, 2014). In our case, we determined what proportion of the distribution of posterior probabilities was different from an expected distribution of values equal to the proportion of females (0.57), adults (0.75), and vocal bats (0.45) in the population (i.e., the subset of individuals that comprised our study, n = 37). Only 6 groups had individuals of both ages (adults and subadults), so we only considered these for the binomial test on age. For the effect of vocal behavior on “dominance” (David’s) scores, we first categorized bats as either vocal (produced at least one response call) or non-vocal (produced no response calls). The beta prior was set to 1, and the results are based on 100000 iterations.

## RESULTS

We collected data on the relative position of 37 individuals belonging to 10 groups of different sizes (two groups of five, three of four, and five of three individuals). We resampled each group 12.5 ± 6.2 times (mean ± standard deviation). Measures of linearity (de Vries’ *h’*) were close to 1, indicating that bats consistently positioned themselves within the tubular leaf in a specific linear order (**Figure 2**). The results of the randomization also show that most individuals (31 out of 37 bats) were consistently selecting the same relative position within the roost (**supplementary figure 1**). The results of the Welch’s ANOVA (F_2,14_ = 3.74, P = 0.05) suggest that linearity varies among positions. The pairwise t-tests show, however, that linearity did not differ significantly among positions (p-values: bottom-middle = 0.11, bottom-top = 0.19, middle-top = 0.33). The results of Levene’s test (F_2,27_ = 4.62, P = 0.01) also show significant differences in the variation of linearity measures among groups within certain positions: the values of all groups for the bottom and top positions are consistently high, whereas there is greater variation in values for the middle position (**Figure 2**). The latter suggests that peripheral positions are more stable while central ones are mostly unstable.

**Figure 2.**
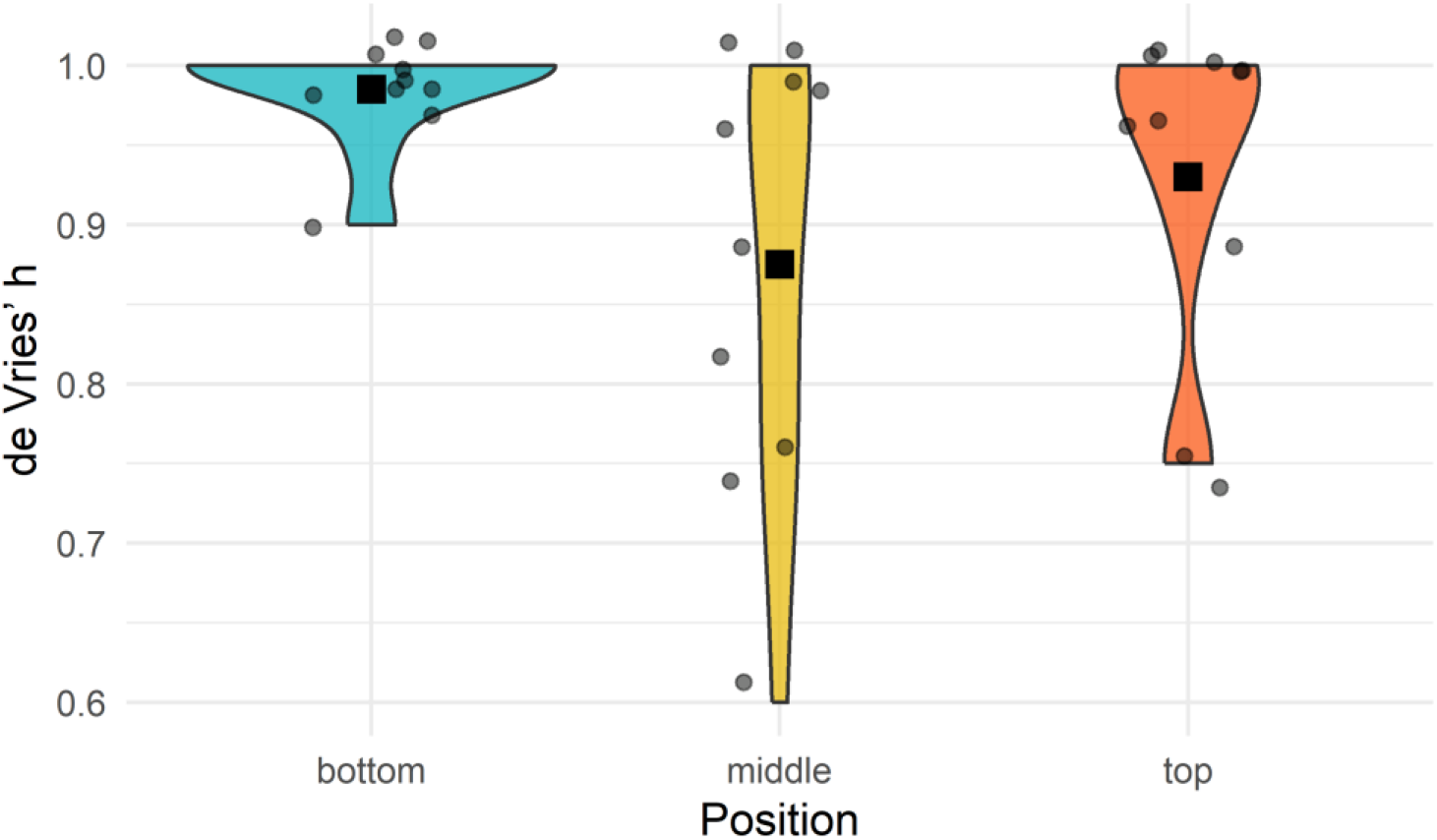
Violin plot of de Vries’ *h’* by the position of dominance, either at the bottom of the roost or at the top (see Figure 1). The black square shows the mean for the position and grey dots show the observed data points.

The dominance index (David’s score) identified which individual(s) consistently occupied a specific position (top, middle, or bottom) within each group (**Supplementary Table 1**). Regarding sex, we found that the relative frequency of success for the top position is more than 0.57, the expected value under the null hypothesis, by a probability of 0.011 and less than 0.57 by a probability of 0.989. This indicates that males have a higher probability of occupying the top position (**Figure 3**). For age, we found that the relative frequency of success is more than 0.75, the expected value under the null hypothesis, by a probability of 0.013 and less than 0.75 by a probability of 0.987. This indicates that subadults are more likely to occupy the top position (**Figure 3**). Contrary to our initial prediction, we found no evidence to suggest that vocal bats were more likely to occupy the bottom position (**Figure 3**).

**Figure 3.**
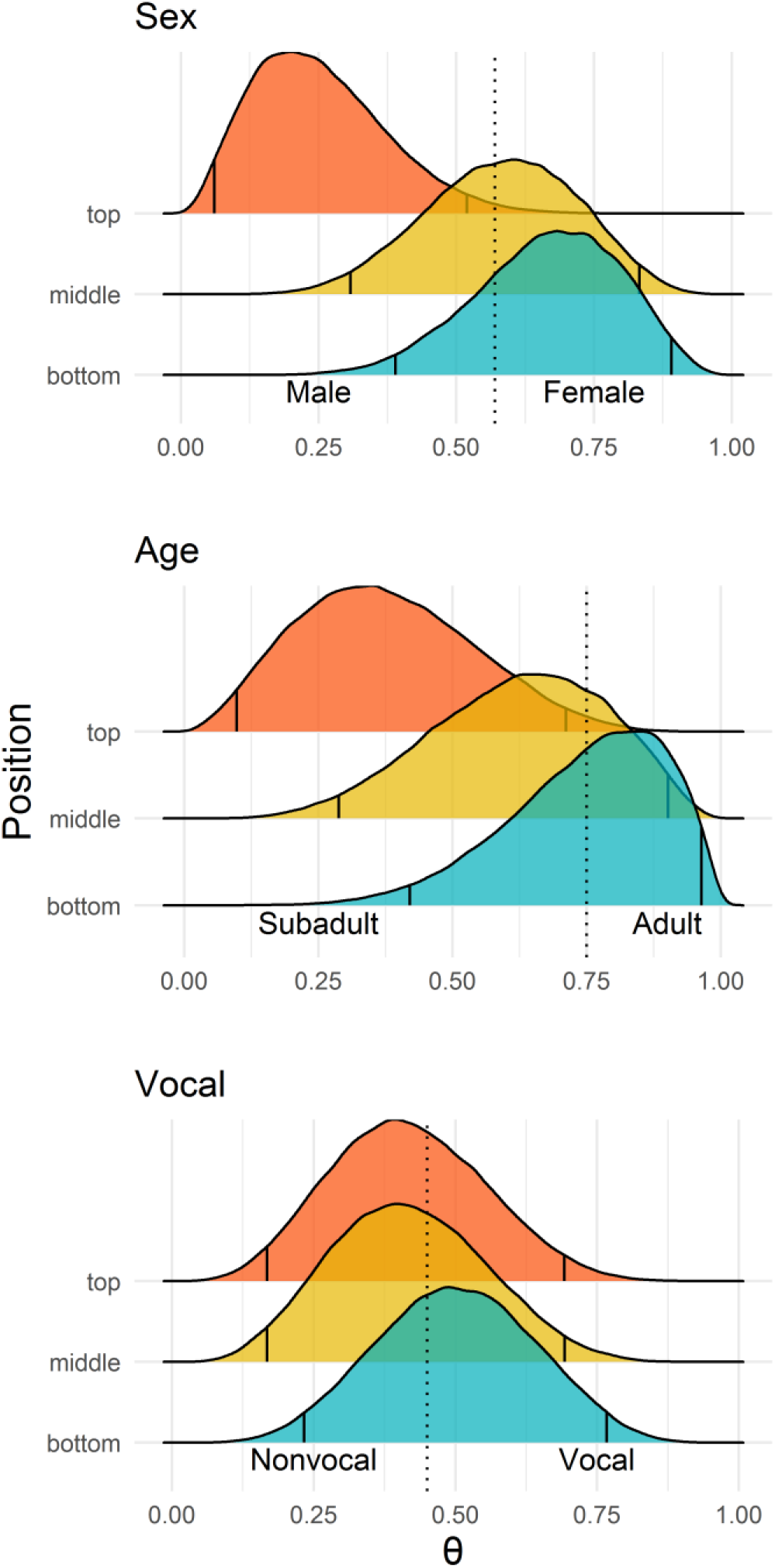
Distribution of posterior probabilities (θ) for the dominance index (David’s score) according to sex, age, and vocal behavior. Dotted lines represent the values expected under the null hypothesis. Vertical lines within the curve indicate the 2.5% and 97.5% tails. Support for the alternative hypothesis, in either direction (male-female, adult-subadult, vocal-non-vocal), is shown whenever the majority of θ values (> 97.5%) fall above or below the dotted line.

## DISCUSSION

In this study we investigated the consistency of individual positioning within the roost in a highly mobile and social bat species (*T. tricolor*). Our results provide the first evidence that bats can consistently position themselves in a predictable manner despite being forced to change roosts daily. We found greater consistency in peripheral positions (top and bottom), while central ones were more labile. Previous studies have also shown that communally-roosting species (e.g., long-tailed tits) exhibit consistent positions within roosts, and that individuals often scramble to gain access to central positions which tend to be more labile than peripheral ones; the latter suggests some form of dominance hierarchy (McGowan et al., 2006). We also found that, when choosing the position inside the roost, there is segregation according to sex and age, with subadult males predominantly occupying the top position. The latter might hint at the presence of variation in resource holding potential or some mechanism that prompts familiar group members to scramble for positions in a predictable manner. Considering the evidence from our study in comparison to others’ (e.g., Summers et al., 1987; Rayor & Uetz, 1990; McGowan et al., 2006), we suggest that future studies could identify if dominance relationships, those in which subordinate individuals submit to dominant ones in contest situations (Kaufmann, 1983), could also influence how bats position themselves within roosts.

High scores for linearity measures within social groups of *T. tricolor* indicate that bats position themselves in a predictable way within the roost, suggesting that relative positioning may depend on 1) relative cost/benefit associated to each position and 2) dominance relationships within the social group. Considering the tubular shape and the verticality of the leaf, the top position might expose bats to the inclemency of the weather, particularly during the rainy season, and to direct attacks of predators that approach the roost from above, such as monkeys and diurnal birds of prey (Boinski & Timm, 1985). Alternatively, being on top may allow bats to escape predators approaching the roost from below, such as snakes (Esbérard & Vrcibradic, 2007). A bat in the bottom position would be protected from the rain but would leave the roost last in the event of a predator’s attack, thus potentially increasing its risk of predation. Considering the pros and cons of the top and bottom positions, we speculate that the middle position is the one that provides the greatest advantages. Central positions have consistently been shown to be more advantageous than peripheral ones (Rayor & Uetz, 1990; McGowan et al., 2006), and our study provides initial evidence that suggests a similar trend for bat groups roosting within developing tubular leaves.

In this study, we found that males and subadults tend to occupy the top position more often than expected by chance. If in fact the middle, or central, position is the preferred one, given the lower consistency in the selection of that location, then our results could support our initial prediction that adults and females have greater RHP and thus relegate subadults and males to the potentially least preferred, riskier, positions (top and bottom). Although we cannot unambiguously interpret our results as outcomes of dominance hierarchy relationships within the group, because we did not investigate antagonistic behaviors between individuals, this positioning pattern is likely to be associated with different social ranks.

We also expected vocal bats to be positioned at the bottom, given that previous studies in *T. tricolor* demonstrate that these individuals are usually the ones to locate roost sites (Sagot et al., 2018). Interestingly, we did not observe a higher probability of vocal bats being at the bottom, suggesting that bats may switch positions once the entire group has entered the roost. The latter would unambiguously demonstrate that positions may not necessarily depend on the timing of arrival, but that individuals scramble to select their preferred position once inside. Further studies are needed to understand which interactions occur within roosts as bats scramble to secure a preferred position.

In conclusion, our results demonstrate that bats of the species *T. tricolor* consistently position themselves inside the roost despite its extreme ephemerality and the high mobility of the species. Whatever the mechanism(s) for the consistent positioning within the roost, this predictable behavior suggests that different positions expose individuals to different costs and benefits. Further research is necessary to understand the latter, and whether dominance relationships represented by antagonistic behaviors influence individual positioning. Typically, the role of roost sites in bats is primarily regarded as promoting affiliative interactions, such as allogrooming, mating, sharing food, and nursing, among others (Kunz, 1982). Our results suggest that roost sites may also serve as venues for establishing relationships based on antagonistic interactions. We also provide initial clues of the formation of dominance hierarchies within social groups, which to date have been rarely identified in bats (Kerth, 2008; Wilkinson et al., 2019).

## ACKNOWLEDGMENTS

We are very grateful to Hal Whitehead for providing advice on estimating linearity and dominance indices in SOCPROG. Annemarie van der Marel, Corina Logan and two anonymous reviewers provided valuable suggestions that helped improve the manuscript. We also thank Julio Bustamante and Lilliana Rubí Jimenez for their help during the research permit application, and Ronald Villalobos for logistics support. This study was also possible thanks to the continuous support of the Centro Biológico Hacienda Barú and its entire staff.

## DATA AVAILABILITY STATEMENT

Analyses reported in this article can be reproduced using the data and code provided in the Figshare repository (https://doi.org/10.6084/m9.figshare.19709485). Additional data may be found in Supplementary Materials.

## AUTHOR CONTRIBUTIONS

Conceptualization: G.G, G.C.; Methodology: G.G, G.C.; Investigation: G.G., S.C.-R., H.A.H.-P., J.P.B.; Formal analysis: G.G, G.C.; Resources: G.C.; Data curation: G.G., G.C.; Writing - original draft: G.G., G.C.; Writing - review & editing: G.G., S.C.-R., H.A.H.-P., J.P.B., G.C.; Visualization: G.C., S.C.-R.; Supervision: G.C.

## ETHICAL STATEMENT

All sampling protocols followed guidelines approved by the American Society of Mammalogists for capture, handling and care of mammals (Sikes et al., 2016) and the ASAB/ABS Guidelines for the use of animals in research. This study was conducted in accordance with the ethical standards for animal welfare of the Costa Rican Ministry of Environment and Energy, Sistema Nacional de Áreas de Conservación, permit no. SINAC-ACOPAC-RES-INV-008-2017 (Decree No. 32553-MINAE). Protocols were also approved by the University of Costa Rica’s Institutional Animal Care and Use Committee (CICUA-42-2018).

## Supplementary materials

**Supplementary Table 1.** Measures of dominance (David’s score) for each individual calculated by group and repeated assuming the top, middle, and bottom position as dominant. Individuals with high scores “dominated” over the others for that position meaning that the individual tended to occupy that specific position (e.g., Gr1-88 dominated when the top position was considered as dominant; therefore, Gr1-88 tended to stay on the top position). Bat ID represents the last 2 digits of the transponder implanted to that individual (for complete transponder data see supplementary figure 1).

| Group – Bat ID | David's score |  |  | Sex | Age | Vocal behavior |
| --- | --- | --- | --- | --- | --- | --- |
|  | Top | Middle | Bottom |  |  |  |
| Gr1 - 22 | -3.00 | -2.75 | -6.00 | Female | Adult | Non-vocal |
| Gr1 - 48 | -3.00 | -2.75 | 3.44 | Female | Adult | Vocal |
| Gr1 - 61 | -3.00 | -2.75 | 7.26 | Female | Adult | Vocal |
| Gr1 - 49 | 4.00 | 4.86 | 1.29 | Male | Subadult | Vocal |
| Gr1 - 88 | 5.00 | 3.39 | -6.00 | Male | Subadult | Vocal |
| Gr2 - 53 | 2.44 | 0.89 | -6.00 | Female | Adult | Vocal |
| Gr2 - 31 | 1.00 | -2.29 | 3.59 | Male | Adult | Non-vocal |
| Gr2 - 95 | -0.50 | -2.29 | 2.36 | Male | Adult | Vocal |
| Gr2 - 75 | -2.94 | 3.68 | 0.06 | Female | Adult | Non-vocal |
| Gr3 - 45 | -3.20 | 2.60 | 2.25 | Female | Adult | Non-vocal |
| Gr3 - 50 | 0.71 | -1.86 | -1.80 | Male | Subadult | Non-vocal |
| Gr3 - 51 | 0.71 | -0.79 | 1.35 | Female | Adult | Vocal |
| Gr3 - 15 | 1.77 | 0.06 | -1.80 | Male | Subadult | Vocal |
| Gr4 - 07 | 3.25 | 3.17 | 3.17 | Female | Adult | Vocal |
| Gr4 - 17 | -3.00 | -6.00 | 3.17 | Female | Adult | Non-vocal |
| Gr4 - 85 | -3.00 | 5.67 | 5.67 | Male | Adult | Vocal |
| Gr4 - 20 | 5.75 | -6.00 | -6.00 | Male | Adult | Non-vocal |
| Gr4 - 34 | -3.00 | 3.17 | -6.00 | Female | Subadult | Vocal |
| Gr6 - 38 | 4.20 | -0.67 | -3.00 | Male | Adult | Non-vocal |
| Gr6 - 78 | -3.00 | -0.67 | 3.80 | Female | Subadult | Vocal |
| Gr6 - 90 | -3.00 | 2.00 | 2.20 | Female | Adult | Vocal |
| Gr6 - 18 | 1.80 | -0.67 | -3.00 | Female | Adult | Non-vocal |
| Gr8 - 33 | -0.97 | 0.22 | 1.17 | Male | Adult | Non-vocal |
| Gr8 - 35 | -0.97 | 0.76 | 0.73 | Female | Adult | Non-vocal |
| Gr8 - 80 | 1.94 | -0.99 | -1.90 | Male | Adult | Non-vocal |
| Gr9 - 15 | 0.33 | 1.00 | 1.38 | Female | Adult | Vocal |
| Gr9 - 19 | 2.67 | -3.00 | -1.75 | Male | Adult | Non-vocal |
| Gr9 - 20 | -3.00 | 2.00 | 0.38 | Female | Adult | Non-vocal |
| Gr10 - 98 | -1.77 | 0.50 | 1.60 | Female | Subadult | Non-vocal |
| Gr10 - 44 | -0.57 | 0.50 | 1.40 | Female | Adult | Non-vocal |
| Gr10 - 66 | 2.34 | -1.00 | -3.00 | Male | Subadult | Vocal |
| Gr11 - 33 | 0.60 | 1.50 | 0.60 | Male | Adult | Non-vocal |
| Gr11 - 95 | 2.40 | -0.75 | -3.00 | Male | Subadult | Non-vocal |
| Gr11 - 29 | -3.00 | -0.75 | 2.40 | Female | Adult | Non-vocal |
| Gr12 - 30 | 2.25 | -1.75 | -1.75 | Female | Adult | Non-vocal |
| Gr12 - 90 | -1.13 | 0.38 | 1.38 | Female | Adult | Vocal |
| Gr12 - 33 | -1.13 | 1.38 | 0.38 | Male | Adult | Vocal |

**Supplementary Figure 1.**
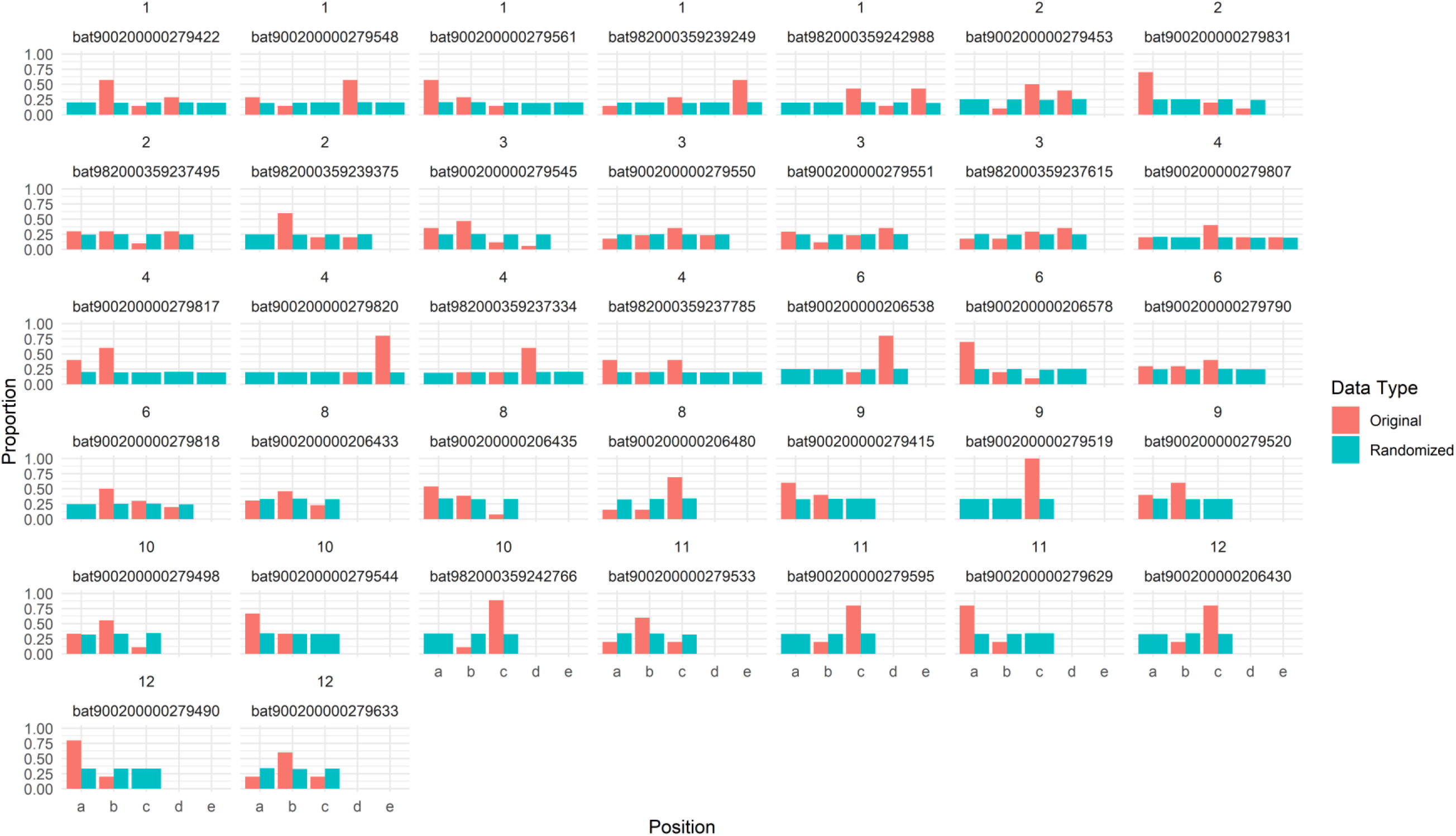
Proportion of times that individuals were found in each relative position within the roost, based on the original or randomized data. The x-axis shows relative position, where *a* denotes the bottom position moving upward towards the roost’s entrance for subsequent letters. Numbers on top of each panel indicate group and bat ID (number of the transponder implanted to each individual).

